# Whole-body 3D kinematics of freely behaving *Drosophila*

**DOI:** 10.64898/2026.05.03.722293

**Authors:** Juan Ignacio Ispizua, Elliott T.T. Abe, Jinyao Yan, Ratan Othayoth, Steven Sawtelle, Felisha Atkins, Hiroshi M. Shiozaki, Niko R. Meier, Jovanka Wong, Tiffany Tran, Catherine Mori, Wilson Chen, Jakob Voigts, David L. Stern, Bingni W. Brunton, John C. Tuthill, Robert Evan Johnson

## Abstract

Understanding how nervous systems generate coordinated movement requires precise measurement of body kinematics during natural behavior. The fruit fly, *Drosophila*, is a model organism with sophisticated behavior and well-studied neural circuits, but tracking fly movements in 3D remains challenging because of their teeny bodies, rapid movements, and frequent self-occlusions. Here we present a pipeline for markerless, full-body 3D pose estimation of fly terrestrial behavior, combining seven synchronized high-speed cameras to capture whole-body kinematics at 800 frames per second. We trained a hybrid 2D/3D deep learning model to track 50 keypoints, then refined them to produce anatomically feasible kinematic trajectories through a retargeting process that solved an inverse kinematics problem constrained by a biomechanical body model. Analysis of 3D kinematics revealed that flies perform grounded running across their full speed range, without transitioning between discrete gaits. Using multi-animal tracking, we found that courting males coordinate both wings during song and modulate body pitch to track the female’s vertical position. Our open-source pipeline and large 3D kinematic dataset of fly behavior provide a foundation for neuromechanical modeling and mechanistic studies of motor control in a genetically tractable model organism.

## 1 Introduction

Quantitative measurement of body movement is critical for understanding the biological mechanisms of animal behavior [1]. Recent advances in deep learning have revolutionized markerless pose estimation, enabling automated tracking of user-defined body parts (i.e., keypoints) across diverse species and behaviors. Two leading software toolkits, DeepLabCut [2] and SLEAP [3], use convolutional neural networks trained on modest numbers of manually annotated frames to achieve human-level accuracy in 2D keypoint detection. Building on these advances, tools such as Anipose [4] and DANNCE [5] have extended pose estimation to 3D, either by combining 2D tracking with multi-camera triangulation or by applying geometric deep learning directly in 3D space [6]. Even with these tools, accurate pose estimation requires high resolution video, and imaging small animals at millimeter scales introduces unique optical challenges. Resolving fine body features demands high magnification, which in turn severely limits depth of field and makes it difficult to keep a freely moving animal in focus as it behaves. As a result, accurate 3D full-body pose estimation in small, fast-moving animals during free behavior has remained out of reach.

The fruit fly, *Drosophila melanogaster*, exemplifies these challenges: at only 2–3 mm in length, flies run at speeds up to 40 mm/s with stepping frequencies exceeding 20 Hz [7]. *Drosophila* has emerged as a powerful model system for investigating how neural circuits control behavior, due to its compact, fully mapped nervous system, advanced genetic toolkit, and rich behavioral repertoire. Flies exhibit sophisticated locomotion and social behaviors that involve complex, dynamically changing body configurations [8, 9, 10]. For instance, during courtship, males perform a probabilistic sequence of motor actions including orientation, chasing, singing, and attempted copulation, each requiring coordinated movements of the legs, wings, head, thorax, and abdomen [11]. Although prior studies have tracked some body parts [12, 13, 14, 15, 16, 17, 18, 19], such as the leg tips, it has not previously been possible to capture the 3D kinematic complexity of behavior in freely behaving flies. Trading off naturalistic behavior for tracking precision, high-resolution 3D joint tracking has previously been achieved only in tethered flies [20, 21], in which the animal’s thorax is fixed while it behaves on an air-suspended spherical treadmill. Tethering substantially alters both the fly’s behavior [22] and locomotor biomechanics, including restricting vertical movements of its body and modifying ground reaction forces.

Accurate 3D kinematics are essential for constraining models of how the nervous system and the body work in closed-loop to generate coordinated movement. Such datasets have been particularly important in training motor control policies capable of simulating biomechanical animal body models to imitate how real animals move [23, 24, 25]. Advances in deep reinforcement learning have enabled efficient training of artificial neural networks to learn such policies [26]. Here again, the fly has been at the forefront of these advances—there now exist two whole-body biomechanical models of *Drosophila* constructed in the open-source MuJoCo physics engine [27, 25]. These models enable researchers to investigate how the biomechanical properties of the body interact with neural control signals to produce realistic behavior, as well as infer sensory inputs to the nervous system that cannot be directly measured (e.g., forces produced by interacting body parts [28] or images of a complex terrain viewed from the retina [27]). The value of neuromechanical models depends on the availability of high-quality kinematic data for training and validation. Existing fly body models have been trained using imitation learning on keypoints tracked in 2D or in 3D from tethered flies [27, 25, 29]. As neuromechanical modeling matures toward incorporating connectome-derived circuit architectures, there is increasing demand for comprehensive, high-fidelity 3D kinematic datasets from freely moving animals performing a range of natural behaviors.

Here we present a pipeline that combines high-speed cameras with custom lighting and uses fast acquisition and annotation software [30, 31] to enable accurate 3D pose estimation of freely behaving flies. Our approach combines multi-camera high-speed video with optimized calibration procedures and deep learning-based keypoint detection to track 50 keypoints across the fly body, capable of achieving a mean tracking error of *<*25 *µ*m. We used these tools to discover how flies vary their 3D joint kinematics across running speeds and to characterize motor patterns of male courtship behavior at high spatial and temporal resolution. We also show how this new 3D kinematics dataset can be used to constrain neuromechanical models of fly behavior. By providing the community with a large, accessible 3D kinematic dataset of diverse terrestrial *Drosophila* behavior, we aim to facilitate mechanistic studies linking neural circuit activity to the biomechanics of natural behavior.

## 2 Results

### 2.1 The rainbow rig tracks whole-body 3D kinematics in freely behaving fruit flies

To measure joint and body kinematics in flies moving freely on the ground, we designed a custom behavioral chamber (24 *×* 5.5 *×* 9.5 mm) constructed from a 3D-printed plastic corridor and two coverslips forming a 30-60-90 triangular prism (**Fig. 1a,b**). Seven high-speed cameras, each equipped with a telecentric lens, were mounted in a semi-circle around the chamber, providing multiple overlapping views of the fly with sufficient depth of field to keep the fly in focus throughout the arena. To capture images at 800 frames per second without motion blur, we illuminated the arena with custom high-power LEDs (**Supplementary Fig. 1**) that strobed synchronously with the camera exposure (50 *µ*s). The short LED duty cycle (4% ON) provided sufficient lighting while limiting hardware heating and perceived brightness. We combined this lighting with GPU-accelerated recording software [30] to enable real-time compression of synchronized video streams. We refer to this behavioral recording apparatus as the “rainbow rig” for its semicircular camera arrangement and color-coded camera views (**Fig. 1c**).

**Figure 1:**
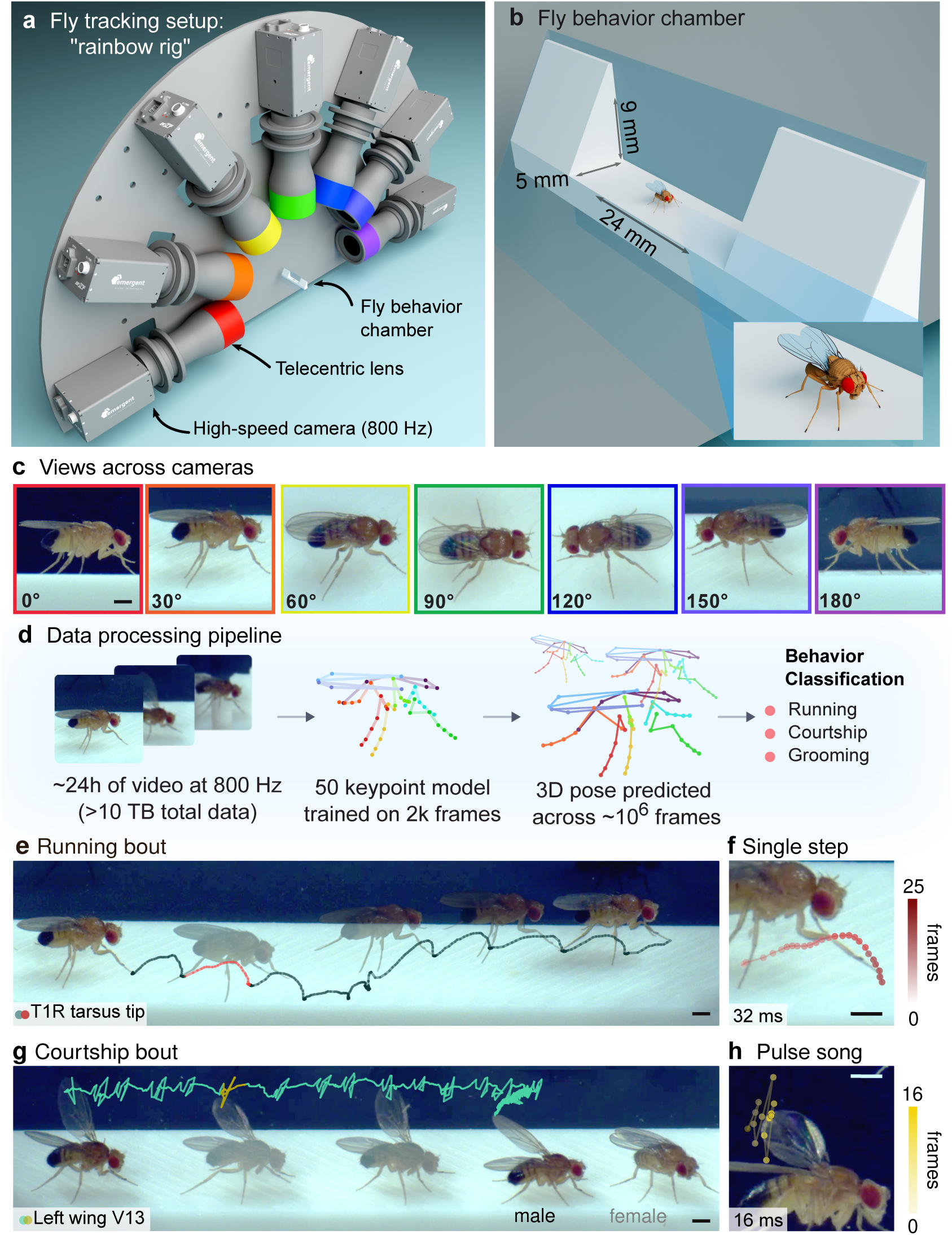
Overview of the behavioral apparatus and 3D tracking pipeline. **a**, Schematic of the rainbow rig for multi-camera high-speed imaging of freely behaving flies. Each high-speed camera is equipped with a telecentric lens focused on the fly chamber. **b**, The 3D-printed fly behavior chamber. The glass walls and floor form a triangular prism enclosure with angles *α* = 60*^◦^* and *β* = 90*^◦^*, such that the upper seam does not occlude the top-down view. To simplify calibration, each camera views the fly through a single plane of glass. **c**, Example frames across all camera views, color-coded to match the cameras in **a**. Multiple views enable precise triangulation of keypoints. **d**, Workflow for generating the dataset of joint kinematics from freely behaving flies. Keypoint labeling was performed in *red* [31], a GPU-accelerated multi-camera annotation application. To train a JARVIS HybridNet, a hybrid 2D/3D neural network for pose estimation, we annotated *∼*2,000 frames per camera view (50 keypoints/frame; the skeleton model includes all leg joints except the middle- and hind-leg coxa-thorax joints). Skeleton overlays show predicted poses. The resulting library of body kinematics was classified into defined behavioral bouts. **e**, Example trajectory of a leg tip for a running bout. **f**, Leg tip trajectory for a single step (red line shown in **e**). **g**, Example wing tip trajectory for a courtship bout. **h**, Wing tip trajectory of a pulse song during courtship inset (yellow line in **g**). All scale bars are 0.5 mm.

To track fly pose from this video data, we developed extensions to JARVIS [32], a hybrid 2D/3D convolutional neural network. Like other deep learning-based markerless tracking tools, JARVIS requires annotating example frames and training a neural network model to predict keypoint locations. To facilitate annotation across multiple views, we used multi-camera annotation software (called *red* [31]) to label a 50-keypoint skeleton in *∼*2,000 frames per camera (*∼*14,000 images total) spanning multiple behaviors. Because *red* uses the camera calibration parameters to triangulate annotations across camera views, only *∼*4,000 images had to be annotated by hand. The skeleton model consists of keypoints on the legs, wings, head, thorax, and abdomen, including all leg joints except the middle- and hind-leg coxa-thorax joints (**Fig. 1d**, **Table 2**). We aggregated annotations across frames, cameras, and behaviors to create a unified JARVIS model for fly pose estimation, then used this model to track keypoints across a large dataset of diverse fly behaviors (43 male and 32 female adult flies, 2–5 days old). Based on patterns of joint kinematics, we then identified and classified distinct fly behaviors, specifically running, grooming, and courtship (**Fig. 1d**, **Supplementary Video 1**).

**Table 1:** Rainbow Rig Hardware Components.

| Device | Vendor & Key Specs | N | Link |
| --- | --- | --- | --- |
| Camera | Emergent Vision Technologies HB-7000-S-C.<br>2.8 MP color camera cropped to $1936 \times 448$ pixels.<br>Recorded at 800 fps with $50 \mu\text{s}$ exposures. | 7 | https://emergentvisiontec.com/products/bolt-hb-25gige-cameras-rdma-area-scan/hb-7000-s/ |
| Lens | $0.36\times$, $1/2''$ GoldTL™ Telecentric Lens.<br>Depth of field: $\pm 3.7$ mm at $f/10$ (20% @ 20 lp/mm) | 7 | https://www.edmundoptics.com/p/036x-half-inch-goldtl-telecentric-lens/15169/ |
| Computer | Puget Rackstation Threadripper PRO WRX90 T131-4U.<br>AMD Ryzen Threadripper Pro 7965WX 4.2 GHz 24 Core 350 W.<br>8× DDR5-5600 ECC Reg. 32 GB.<br>NVIDIA RTX 4000 Ada Generation 20 GB PCI-E.<br>Ubuntu 22.04 LTS w/ Gnome Desktop Installation (64-bit). | 1 | https://www.pugetsystems.com/products/rackmount-workstations/amd-rackstations/t131-4u/ |
| GPU | NVIDIA A16 | 2 | https://www.nvidia.com/en-us/data-center/products/a16-gpu/ |
| NIC | NVIDIA 10/25 GigE Quad-Port 10/25 GigE SFP28.<br>MCX713104AS-ADAT. | 2 | https://emergentvisiontec.com/interface-cards/third-party-adapters/25gige-nvidia-mcx713104as-adat-network-interface-card/ |
| Lighting | Custom Janelia Saturn IV Strobe LED | 2 | See Lighting Section |

**Table 2:**
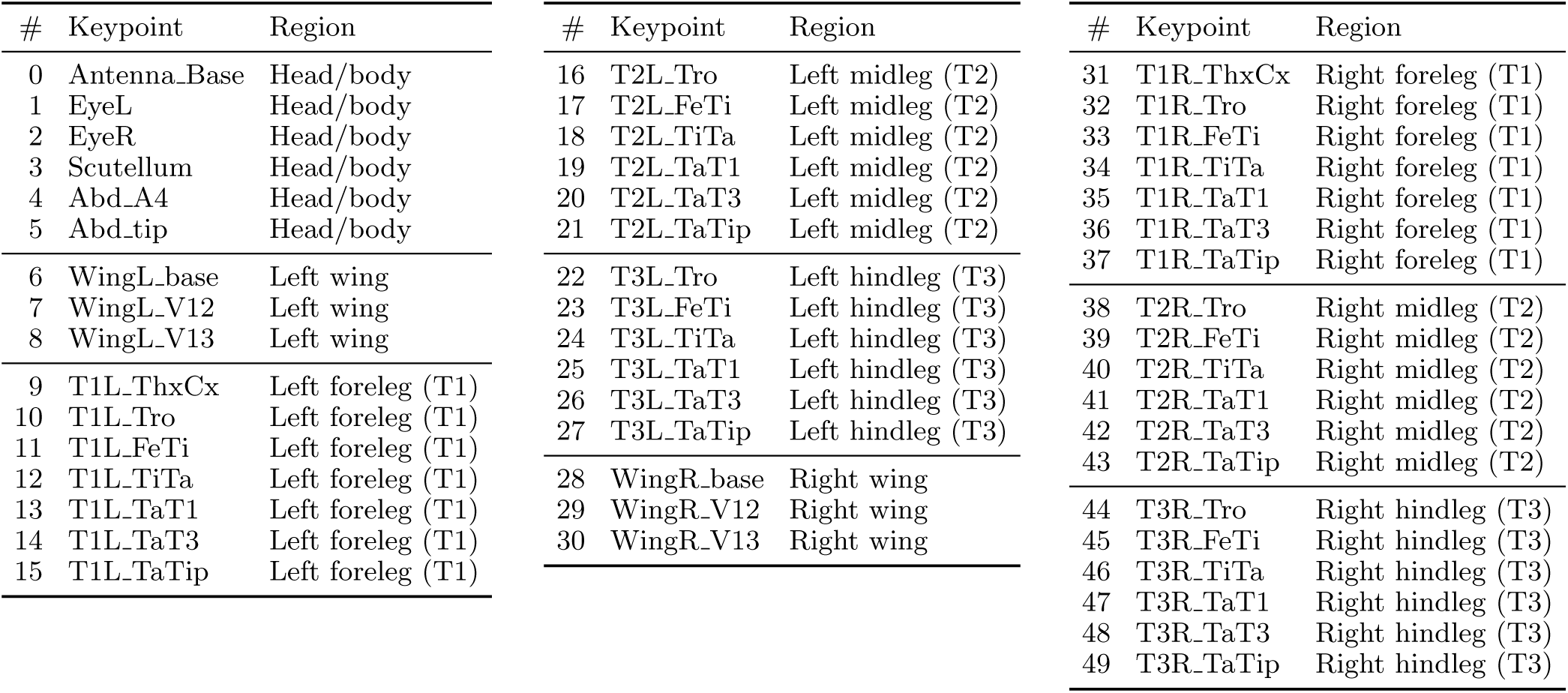
The 50 keypoints of the fly skeleton used for 3D annotation.

The high temporal resolution of the dataset (800 Hz) allowed us to densely reconstruct the trajectory of individual steps during running (**Fig. 1e,f**) and single wing pulses during courtship song (**Fig. 1g,h**). All cameras had *∼*12.3 *µ*m/pixel spatial resolution (1936 x 448 pixels) and viewed the entire arena floor. To evaluate the tracking error of our pipeline, we computed a reprojection error of 0.48 *±* 0.26 pixels (5.9 *±* 3.2 *µ*m; mean *±* s.d.) on 3D calibration landmarks (see Methods). To estimate tracking accuracy on held-out behavioral data, we computed the reprojection error of triangulated 3D predictions for a leg keypoint. Specifically, we manually annotated foreleg tip positions from a running bout across all seven camera views in 51 frames in which the leg tip was visible from all cameras. This analysis obtained an overall keypoint tracking error of 1.48 *±* 0.69 pixels (18.2 *±* 8.5 *µ*m; mean *±* s.d.). For context, this error is approximately 2–3% of the length of a *Drosophila* femur (*∼*0.72 mm [33]).

To refine raw keypoints and produce biomechanically feasible kinematic trajectories for further analyses, we retargeted the predicted keypoints by solving an inverse kinematics problem constrained by a fly body model using STAC (Simultaneous Tracking and Calibration [34, 26]). Like most other markerless pose estimation methods, keypoints predicted by JARVIS have several properties that complicate direct kinematic analysis. Individual keypoint positions were tracked independently so were subject to jitter in adjacent frames from prediction error. Keypoints also do not respect fixed body size and anatomical constraints, such as limb segment lengths and joint ranges of motion. Moreover, estimated keypoint positions report surface landmarks, rather than the underlying joint angles that are under more direct control of the nervous and musculoskeletal systems. STAC refines tracked keypoints by solving an inverse kinematics problem, finding the joint configuration most consistent with the estimated keypoints while enforcing anatomical constraints from a fly biomechanical body model implemented in MuJoCo [25], thus yielding smooth, physically plausible joint angle trajectories in an egocentric reference frame. This registration process was computed for tracked keypoint trajectories over each bout, and smoothness was promoted by temporal regularization. Unless otherwise noted, all kinematic analyses presented below are performed on STAC-refined joint angles, rather than raw keypoint positions.

### 2.2 3D joint kinematics of freely running flies

Freely running flies have been tracked in 2D and tethered flies have been tracked in 3D, but precise and comprehensive 3D joint tracking during free behavior is needed to understand how flies coordinate their legs during adaptive locomotion and how this coordination is controlled by the fly nervous system. Therefore, we first analyzed the leg joint kinematics of running flies (**Supplementary Video 1**). We defined a step cycle as one complete oscillation of the left foreleg (L1) from swing onset to swing onset, equivalent to one stride in bipedal locomotion [35, 36]. We then identified running bouts containing at least two complete step cycles, during which all six legs passed through two full swing and stance phases (**Fig. 2a**). This criterion resulted in a curated dataset of 2213 running bouts (*∼*654,000 frame-sets) across various speeds, body heights and headings (**Supplementary Fig. 2a-d**). From the smooth joint angle traces in these bouts (**Fig. 2b**) we could extract comprehensive joint angle distributions (**Fig. 2c**) across all six legs.

**Figure 2:**
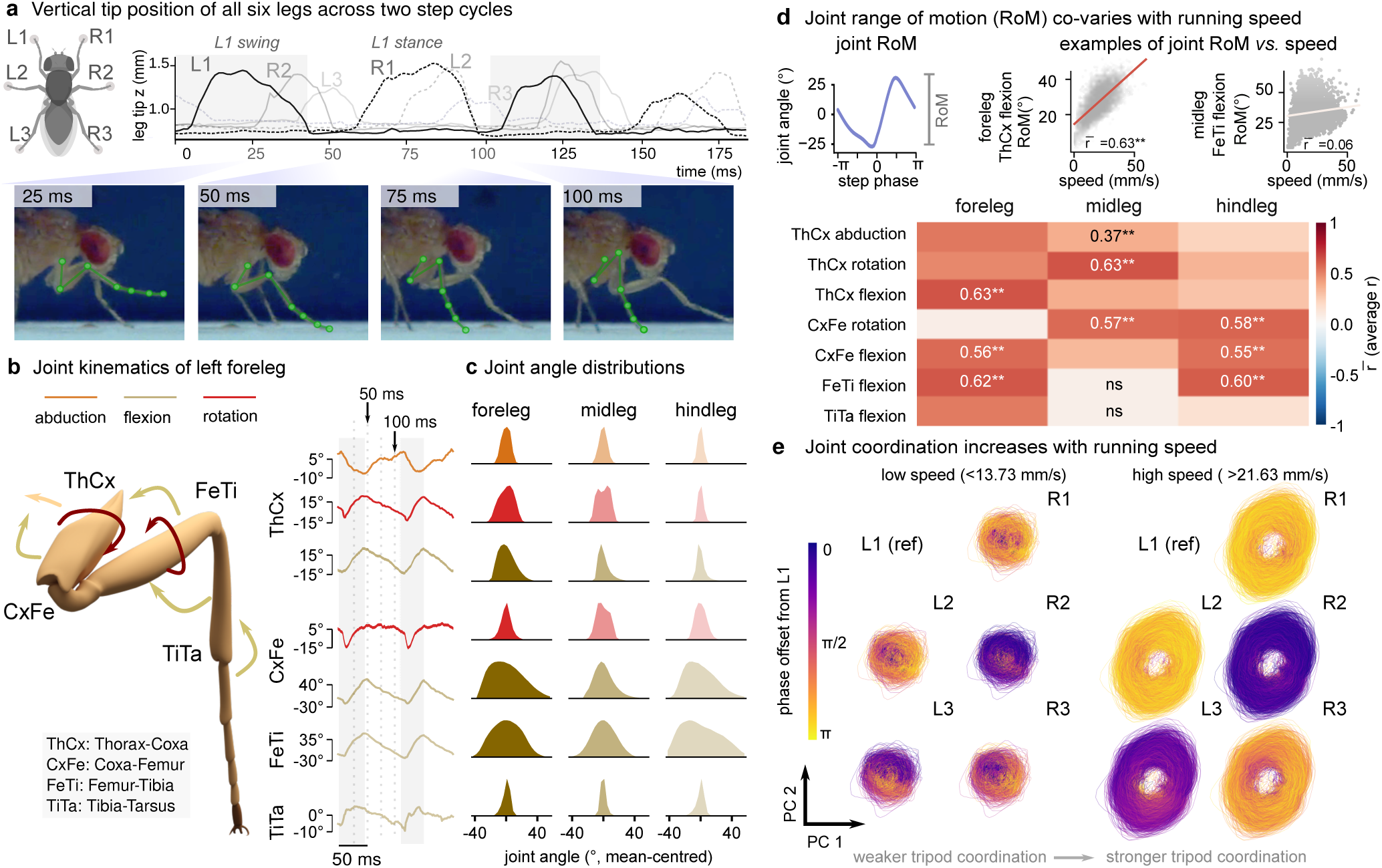
3D leg joint kinematics of freely running *Drosophila*. **a**, Vertical leg tip position across two step cycles. Images below highlight different left foreleg (L1) poses across the step. Although the leg tip does not change position during stance, other leg joints move through a trajectory of angles. **b**, Schematic of a foreleg showing the 3D angles calculated through inverse kinematics (left). At right are individual joint angle traces for the L1 leg. The dotted lines correspond to the inset images in **a**. **c**, Leg joint angle distributions across all running bouts in the whole dataset (53 flies, 2213 running bouts, averaged across symmetric pairs of legs). All distributions are normalized to their maximum value. Different joints have different ranges of motion (RoM). **d**, Co-variance of joint RoM with running speed. The RoM of each joint was calculated by taking the difference between the maximum and minimum angle values per cycle, averaged bilaterally across pairs of legs. A linear regression was calculated per fly and then averaged across the dataset to produce the RoM vs. speed correlation matrix. Only the three joint angles with the highest correlation per leg are shown for simplicity. **e**, Principal component analysis (PCA) of all leg joint angles and their derivatives. Trajectories of first two principal components are shown. For each leg, trajectories are colored by circular phase offset relative to L1, showing differences across legs in coordination and stereotypy of running at high and low speed.

To understand how flies adapt their joint kinematics during locomotion at different speeds, we characterized how the range of motion (RoM) of each joint angle changed as a function of running speed (**Fig. 2d**). While the RoM of some joints increased monotonically with speed, others remained unchanged, and these speed-dependent differences were not uniform across legs. In the front and hind legs, RoM increased with speed primarily at the femur-tibia and tibia-tarsus joints, whereas in the middle legs, speed-dependent increases were concentrated at the femur and coxa rotation joints. This observation about how the middle vs. front/hind legs adjust to higher speeds is consistent with previously described differences in the mechanics of the middle legs relative to the front and hind legs, which are thought to play distinct roles in steering and propulsion during insect locomotion [37, 38, 13].

To visualize how inter-limb coordination changes across speeds, we performed principal component analysis (PCA) on all leg joint angles and their angular velocities across all six legs. Because running is highly rhythmic, the first two principal components (PCs) captured the dominant oscillatory structure of locomotion, and plotting kinematic trajectories through PC1-PC2 space yielded closed circular orbits corresponding to individual step cycles (**Fig. 2e**). We next computed the phase offset of each leg relative to the left foreleg (L1). Coloring stepping trajectories by this phase offset revealed how inter-limb coordination changed with speed. At high speeds, legs belonging to the canonical tripod groups (L1/R2/L3 and R1/L2/R3) shared similar phase offsets, reflecting stereotyped tripod coordination [15]. At low speeds, trajectories became less regular and phase offsets were more variable; in particular, the hind legs showed weaker coupling to L1 (**Supplementary Fig. 2e-f**). This degradation of tripod structure at low speeds is consistent with results from 2D leg tip tracking studies in flies [18, 19], and our results extended those findings by demonstrating that the change in coordination strength is distributed across the full joint angle space, rather than being restricted to any single joint or leg segment.

### 2.3 Fly kinematics are consistent with grounded running at all speeds

Walking and running are not simply faster and slower versions of the same locomotor pattern; they are kinematically and dynamically distinct, suggesting they engage distinct mechanical strategies with different implications for neural control [36, 35]. In legged animals, one way walking and running are distinguished is by the phase relationship between fluctuations in kinetic and potential energy across the step cycle [35]. During walking, these quantities oscillate out of phase, as in an inverted pendulum, whereas during running, they oscillate in phase, as the animal decelerates and loads a compliant leg spring before rebounding. Consistent with a cyclic storage and exchange of elastic energy into kinetic energy, the body center of mass (CoM) *height* and *speed* also vary out of phase for walking and in phase for running (**Fig. 3a**). Prior work used 2D tracking of fly locomotion to show that flies use a modified tripod coordination pattern across all locomotor speeds [18, 17]. Further, an estimated CoM tracked from a side view was used to resolve that the geometry of the tripod, rather than individual leg stiffness, determines the effective leg spring stiffness [17]. However, without full 3D tracking of body and leg joint kinematics in freely behaving flies, the phase relationship between CoM height and speed across the step cycle could not be directly measured.

**Figure 3:**
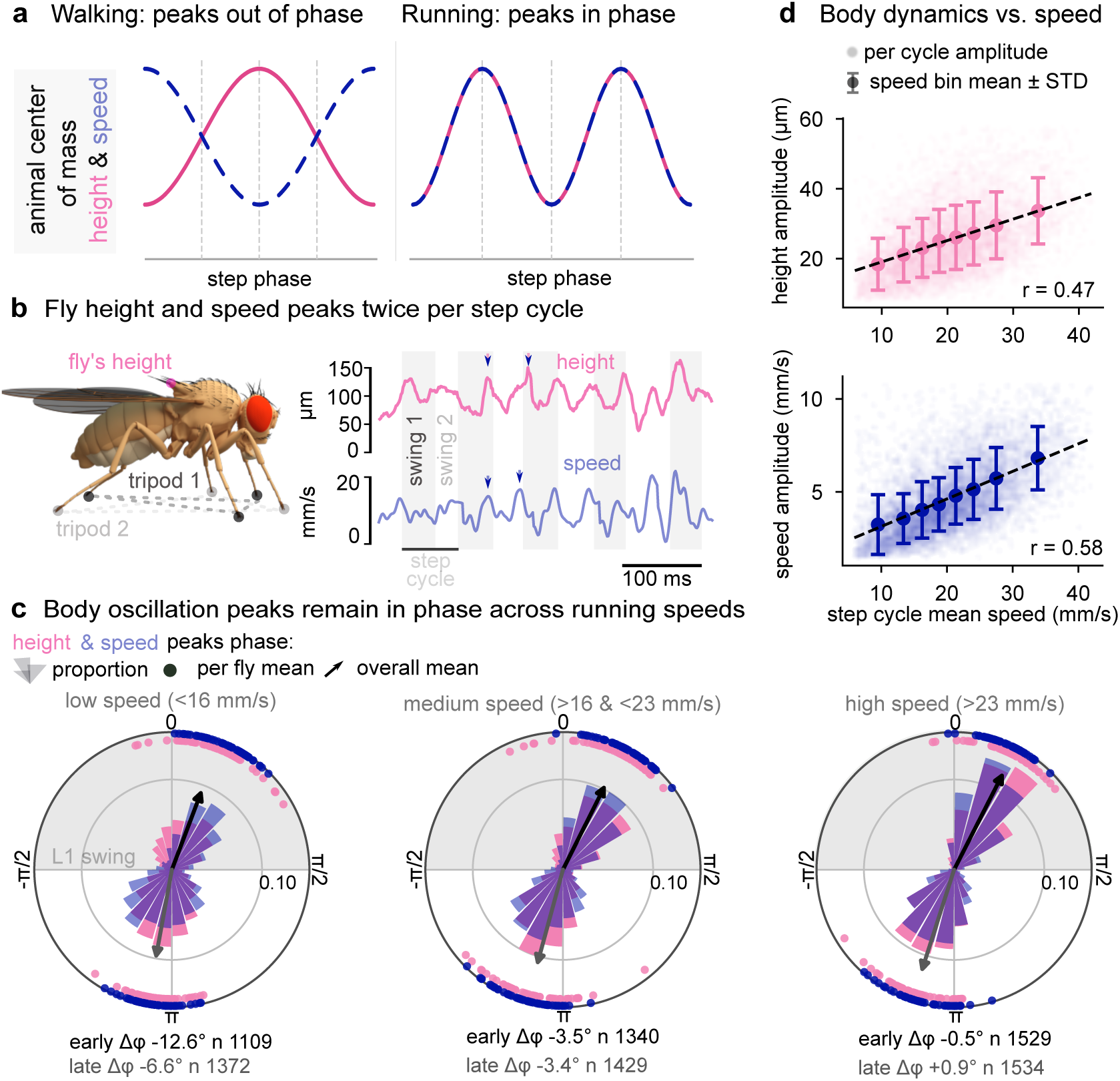
Fly locomotor dynamics resemble running more than walking. **a**, Schematic description of how body center of mass (CoM) dynamics vary with step phase during walking and running. According to the spring-loaded inverted pendulum (SLIP) biomechanical model of legged locomotion, running and walking are distinguished by the peaks of these quantities fluctuating in and out of phase, respectively. **b**, Example traces of fly height and speed across multiple step cycles. Both quantities peaked during the swing phase of each alternating tripod. Height was detrended to remove slow changes in posture by subtracting the mean across a sliding 100 ms window. **c**, Polar plots showing the distribution of the height and speed peaks as a function of step phase, using the L1 leg as reference. Polar histograms show the probability densities, as well as mean phase per fly (circles) and across flies (arrow), of the first and second peaks of the step cycle (noted as *early* and *late*). Both the mean phase and distribution width remain fairly unchanged across speed tertiles. Phase difference is noted as Δ*ϕ* . **d**, Height and speed oscillations, calculated as the amplitude of their fluctuations about the detrended per-cycle means, are correlated with the mean speed per step cycle. Both the binned mean values and linear regression are shown. Slopes of linear fit lines are 0.159 and 0.567 for height and speed, respectively.

Using speed and height changes at the scutellum as a proxy for those at the CoM of the fly’s rigid body, we found that both quantities peak twice per step cycle (i.e., twice per stride in standard bipedal terminology), phase-locked to the swing phases of the two alternating tripods (**Fig. 3b**). This coordination pattern is analogous to the two diagonal limb pairs in a trotting quadruped [39, 40]. The first peak falls between 0.5 and 0.6 radians (mid L1 swing) and the second between *−*2.8 and *−*2.9 radians (mid L1 stance). Critically, height and speed peaked in phase, at both alternating tripod peaks, across the full range of locomotor speeds (**Fig. 3c**). This in-phase relationship is one defining signature of running rather than walking mechanics [35], although these running-like kinematics in flies did not include a “floating” phase where all leg tips are suspended off the ground, as is the case in quadruped running. These results indicate that *Drosophila* locomotion is biomechanically more analogous to running than to walking at all speeds, in a coordination pattern consistent with *grounded running* or *Groucho running* [41, 42]. The amplitude of both oscillations scaled with mean speed (**Fig. 3d**), but their phase relationship remained stable, consistent with the interpretation that flies do not transition between discrete gaits but instead modulate a single running-like locomotor program continuously as a function of speed.

### 2.4 Multi-animal tracking of flies during courtship

Using the rainbow rig, we recorded social interactions between male and female flies (11 pairs). We then trained a custom multi-animal model (following a similar approach to [43]) to track both animals during courtship (see example in **Fig. 4a** and **Supplementary Video** 2). Male flies sing in the presence of females, extending and vibrating one of their wings. While past work has used microphones to record fly courtship songs [44, 45, 46, 47], the high temporal resolution of our video data allowed us to measure courtship song directly from wing kinematics. In *Drosophila melanogaster*, courtship song consists of two stereotyped syllable types: *pulse* song, characterized by short, high-amplitude wing oscillations separated by regular intervals, and *sine* song, a sustained low-amplitude oscillation produced when the male is in close proximity to the female [48, 49, 50]. Both song types were readily identifiable in our recordings from the characteristic trajectory of the wing tip (**Fig. 4c,e**), and we were further able to distinguish two subtypes of pulse song based on differences in waveform shape (**Fig. 4f**).

**Figure 4:**
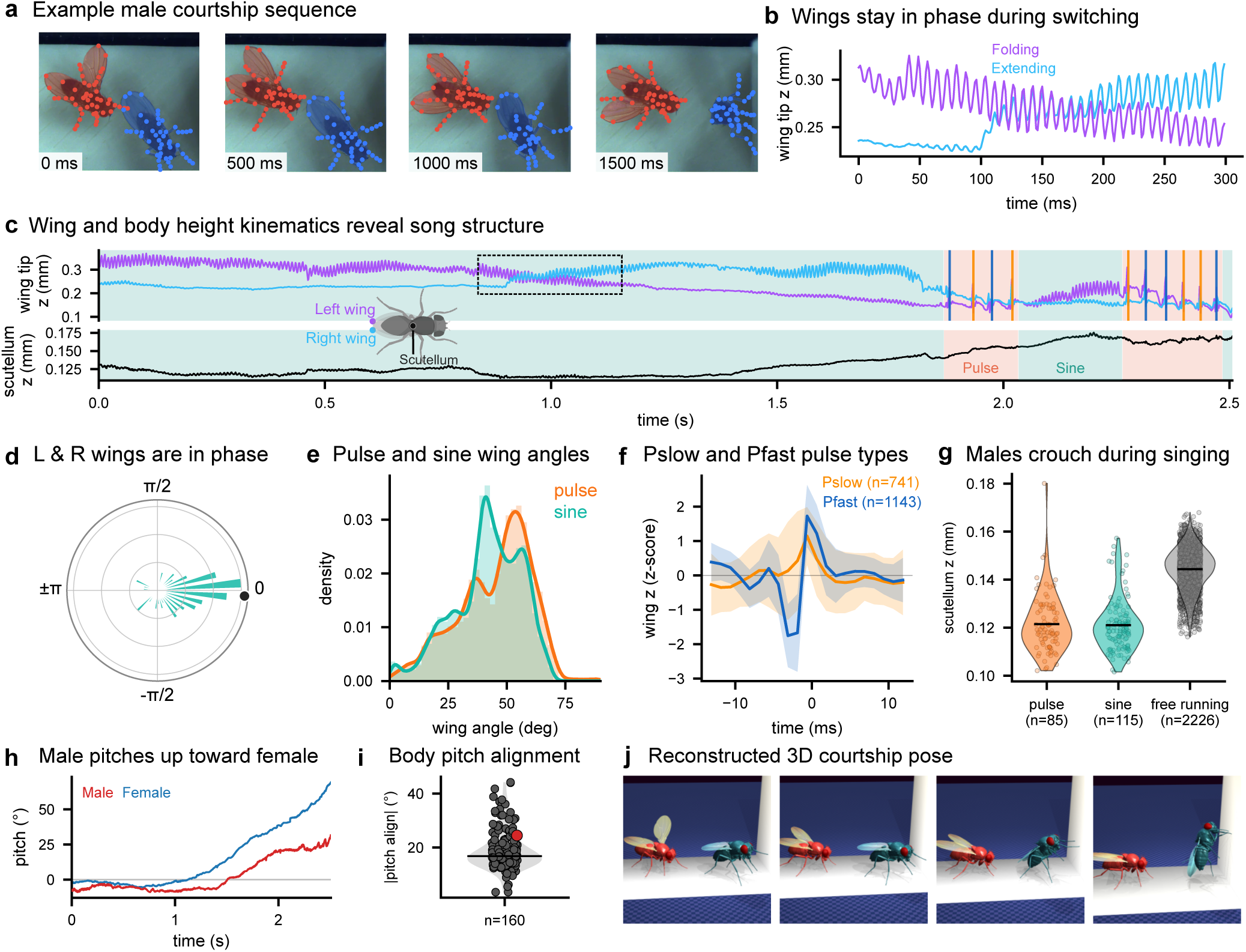
3D tracking of multi-animal social behavior during courtship. **a**, Example video frames sampled across the bout, overlaid with projected keypoints (red = male, blue = female) and per-animal segmentation masks, demonstrating multi-animal detection and pose estimation. **b**, Example of sine song switching, illustrating that the wings oscillate in-phase throughout sine song. This example trace is indicated with a dashed box in **c**. **c**, Wing tip *z*-position (L, purple; R, blue) over the exemplar bout, with scutellum *z* stacked below. Pulse and sine song segments are shaded behind the traces (salmon = pulse, teal = sine); vertical lines mark individual pulse events colored by sub-type (*P*_slow_ = orange, *P*_fast_ = deep blue). **d**, Polar histogram of the phase difference between wings for all sine songs; they are concentrated near 0 rad, confirming consistent wing-phase coupling. Black dot is median phase difference. **e**, 2D density of the absolute value of horizontal wing angle (defined as the angle between the body axis and the horizontal projection to the wing tip on the fly body model) during pulse and sine songs. An angle of zero represents the wing’s resting position (folded on the back). *N* =11 flies. **f**, Pulse-waveform classification: mean *±* SD of pooled *P*_slow_ and *P*_fast_ waveforms classified by a two-component gaussian-mixtures model, pooled across all pairs. **g**, Scutellum *z*-height distribution during singing bouts vs. free running, showing the postural elevation associated with courtship. **h**, Male (red) and female (blue) thorax pitch relative to the floor throughout a courtship bout. **i**, Per-bout median of the angle alignment between male singing and female center of mass. The exemplar bout shown in **j** is highlighted in red. **j**, MuJoCo-rendered poses spanning the same time range as **c**.

Sine song is generally thought to be produced by vibration of one extended wing, whereas the contralateral wing is not extended and does not contribute to song [51, 52]. Our high-resolution wing tracking revealed that the non-extended wing nevertheless oscillates in phase with the singing wing during sine song (**Fig. 4d**). This bilateral coupling was most evident during wing switches, in which the amplitude of the previously extended wing gradually decreased while that of the opposite wing increased, with no discontinuity in oscillation phase across the transition (**Fig. 4b,c**). The preservation of phase across the switch suggests that the indirect flight muscles, which are known to power both flight and courtship song through bilateral deformation of the thorax [52, 53], sustain continuous bilateral contraction throughout the transition. The asymmetry in wing amplitude between the singing and silent wings may, therefore, arise not from asymmetric indirect muscle activation, but from active neural suppression of the non-singing wing [52]. This finding illustrates how high-resolution video-based pose estimation can reveal aspects of courtship motor control that were not previously identified through microphone-based recordings.

The high temporal and spatial resolution of our recordings also enabled quantification of whole-body postures during courtship. We found that males adopted a significantly lower body posture during singing, for both pulse and sine song, compared to free running (**Fig. 4g**). Beyond body height, our 3D tracking revealed that males actively modulated their pitch angle relative to the female’s position: the male’s thorax pitch closely tracked the angle between his body and the female’s center of mass (**Fig. 4h**), with a median angular offset of 23.2*^◦^* across bouts (**Fig. 4i**). This pitch alignment, which could not be resolved from 2D recordings, suggests that males orient not only in the horizontal plane as previously described [50], but also in the vertical plane relative to the female. This 3D body orientation may influence how courtship song is directed toward and perceived by the female [54]. Finally, multi-animal inverse kinematics allowed us to reconstruct the full 3D pose of both animals simultaneously throughout each courtship bout (**Fig. 4j**, **Supplementary Video 3**), providing a rich kinematic representation of social interactions that could be used to directly constrain neuromechanical models of coordinated multi-animal behavior, including potential reconstruction of tactile interactions.

### 2.5 3D tracking of other behaviors and experimental conditions

To demonstrate the versatility of the rainbow rig and 3D tracking pipeline, we recorded and tracked several fly behaviors beyond running and courtship. We first focused on grooming, which flies performed spontaneously within the chamber. Grooming is a particularly demanding behavior to track because it involves rapid, complex movements of multiple body parts in close proximity and contact, with frequent occlusions between the legs and head [55]. During anterior grooming, for example, flies alternate distinct foreleg movements: sweeps across the antenna, head and eyes, and bilateral rubbing of the forelegs together to remove debris. These mutually exclusive actions must be resolved at the keypoint level [56]. Despite these challenges, we found that augmenting our training data with *∼*600 additional annotated frames was sufficient to achieve accurate tracking of head and foreleg keypoints throughout several anterior grooming bouts, allowing us to resolve distinct grooming syllables and to document the full range of motion of the head (**Fig. 5a**, **Supplementary Video 4**). We also found that the midlegs shift their stance outward and anteriorly, which may improve postural stability during foreleg lifting (**Fig. 5b**). We also demonstrate the capacity of the rainbow rig to record full 3D body kinematics during optogenetic manipulations. We expressed CsChrimson in DNg100, a descending neuron previously identified as a command neuron sufficient to drive rhythmic running in headless flies [57, 58]. Illuminating the arena with a red LED reliably elicited sustained running bouts, which we tracked with a modified headless skeleton (**Fig. 5c**, **Supplementary Video 5**). The resulting 3D joint kinematics provide a baseline for comparing the kinematic signatures of descending command neuron activation with those of spontaneous, brain-driven locomotion. We found that even though induced running uses a narrower range of leg flexion, its kinematics are still a stereotyped subset of wild-type running in the principal component space (**Fig. 5c-d**). Finally, we demonstrate 3D pose estimation of flies navigating complex terrain. We designed and 3D-printed a series of steps within the behavioral chamber, challenging flies to traverse vertical obstacles during spontaneous locomotion. Flies dexterously negotiated the steps (**Fig. 5e**, **Supplementary Video 6**), frequently pausing at step edges to perform exploratory foreleg searching movements and performing double leg swings after failing to contact the ground during a running bout (**Fig. 5f**). These movements were reminiscent of those previously described during gap-crossing, in which flies use visual and tactile cues to assess and reach across chasms [59].

**Figure 5:**
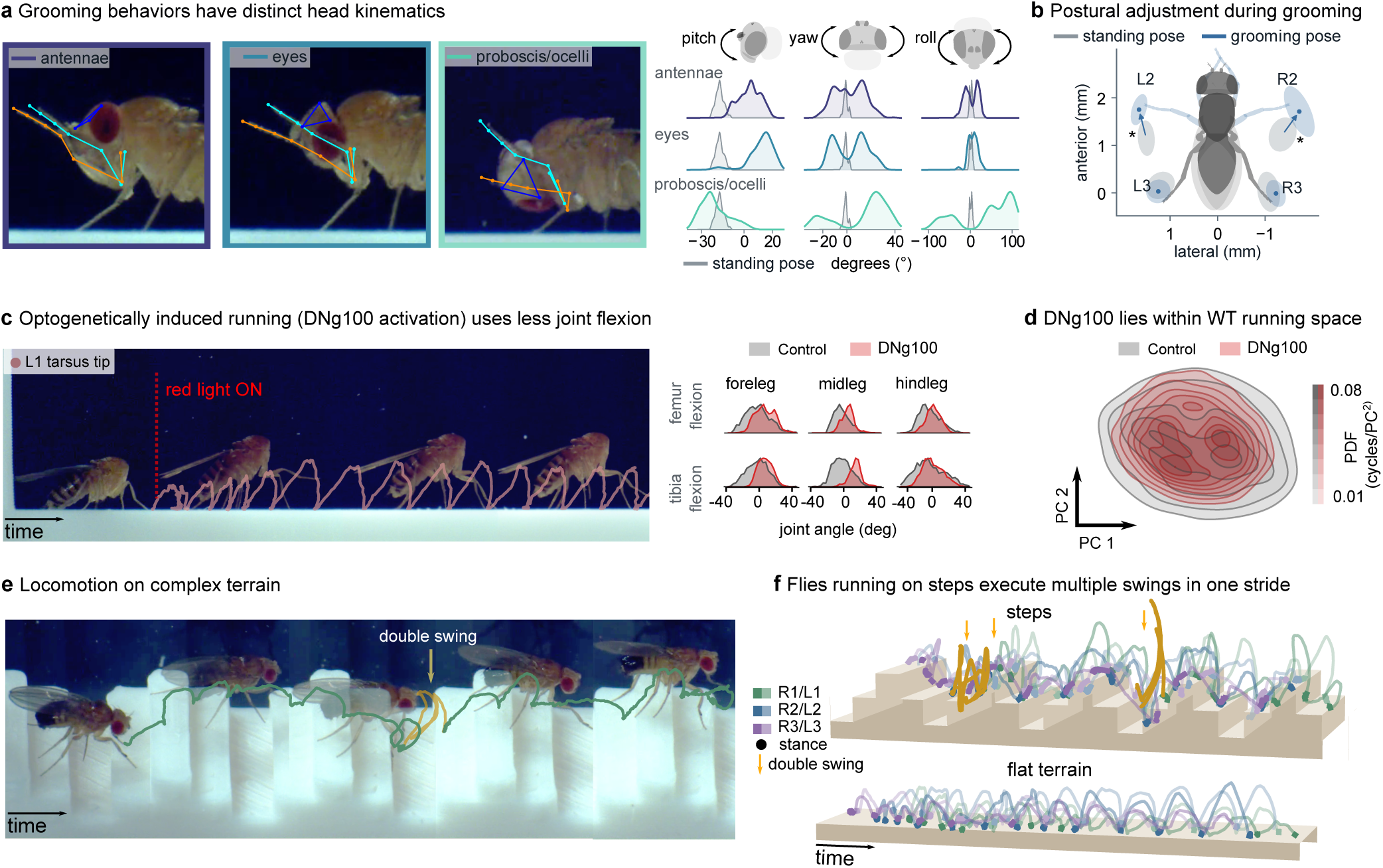
3D tracking of other behaviors and experimental conditions. **a**, Example images of a fly performing 3 distinct anterior grooming patterns (antennal grooming, eye cleaning and proboscis/ocellar grooming). Head angle distributions (pitch, yaw and roll) are shown for different grooming behaviors, compared to standing pose. **b**, Midlegs shift position during grooming. Ellipses are the 50% probability contours of leg-tip XY position. Centroid shifts 0.41 mm for L2 and 0.47 mm for R2, statistically significant (p = 0.002, two-sample Hotelling *T* ^2^, Holm-Bonferroni corrected for multiple comparisons over six legs). n=14 grooming bouts (16,072 frames), 19 standing bouts (2,123 frames), same male fly. Individual frames are from **Supplementary Video 4**. **c**, Tracking running behavior of optogenetically induced locomotion (DNg100*>*CsChrimson) in a decapitated fly. Composite image constructed from **Supplementary Video 5**. Flexion at the tibia and femur is reduced during induced running. Joint angle distributions are mean centered on the wild-type condition and normalized to their maximum value. **d**, Probability density in the plane of the first two principal components (PC) for DNg100 and wild-type speed-matched running cycles. Step cycles of the experimental conditions were projected onto the full wild-type PC space. The centroids of both distributions are only 0.17 PC units apart, and the experimental condition is fully contained on the wild-type space. n=50 running cycles per condition, speed matched. **e**, Tracking 3D body kinematics of a fly running on complex terrain; composite image from **Supplementary Video 6**. **f**, 3D leg-tip trajectories of two running bouts, one on complex terrain (’steps’) and one on even terrain, demonstrating changes in the leg swing shape and height, as well as episodes where legs repeated swing after a failed landing.

Accurate tracking of flies during these additional behaviors required only a small number of additional annotated frames, further illustrating the efficiency with which our pipeline can be adapted to new behavioral contexts. These recordings provide the first 3D joint kinematic dataset of *Drosophila* navigating structured terrain, opening the door to quantitative analysis of how flies modulate leg trajectories, body posture, and inter-leg coordination in response to environmental challenges.

## 3 Discussion

In this paper, we introduce a new integrated pipeline for high-fidelity 3D pose estimation in freely behaving *Drosophila*. By combining new optical hardware with pose detection and robust triangulation algorithms, we overcame the unique challenges that had previously prevented tracking 3D kinematics of small, fast-moving arthropods like flies. We demonstrate the capacity of our setup to accurately track 50 keypoints distributed across the fly’s body, including legs, wings, and head. These data enabled the first comprehensive 3D kinematic analysis of freely moving *Drosophila*, including locomotion and wing dynamics during courtship. This work provides a foundation for mechanistic studies linking neural circuits to the biomechanics of natural behavior in this well-studied model organism.

### 3.1 3D kinematics reveal adaptive motor strategies during locomotion

Our analysis of running kinematics revealed how *Drosophila* coordinate 3D joint configurations across a continuous range of locomotor speeds. Rather than transitioning between discrete gaits, flies modulate a single locomotor program continuously, with speed-dependent changes in joint range of motion. These changes are distributed non-uniformly across leg pairs: femur-tibia and tibia-tarsus joints in the front and hind legs, and femur and coxa rotation joints in the middle legs. PCA of the full joint angle space confirmed that inter-limb coordination becomes more stereotyped at higher speeds, with canonical tripod groupings emerging clearly in the phase structure of low-dimensional trajectories, while slow running is characterized by weaker and more variable coupling, particularly in the hind legs. This speed-dependent change in coordination suggests that the nervous system shifts from flexible, feedback-dependent control at low speeds to more rigid, centrally-patterned coordination at high speeds, consistent with a transition from sensory-driven to CPG-dominated locomotor control.

Analysis of body dynamics revealed that both height and forward speed peak in phase with each other across the full speed range. This phase relationship is one defining signature of running rather than walking biomechanics [36, 35], indicating that *Drosophila* locomotion is better described as using running-like spring-mass dynamics at all speeds. This finding was made possible using 3D pose estimation of freely behaving flies and provides a useful constraint for neuromechanical models of fly locomotion.

### 3.2 Courtship kinematics in three dimensions

Our multi-animal tracking of courtship behavior demonstrated that high-resolution 3D pose estimation can reveal novel aspects of motor control and social behavior. By measuring wing tip kinematics at 800 Hz, we identified both pulse and sine song directly from wing trajectories. We also distinguished two subtypes of pulse song based on waveform shape, consistent with the P*slow* and P*fast* classification established from acoustic recordings [44]. We found that during sine song, both wings move and oscillate in phase, including the wing conventionally considered to be silent [51, 52], although whether oscillations of the non-extended wing produce any sound remains unknown. Bilateral phase coupling was maintained without interruption during wing switches, with oscillation amplitude transferring smoothly from one wing to the other and no discontinuity in phase across the transition. This kinematic signature is consistent with continuous bilateral contraction of the indirect flight muscles throughout the switch [52, 53], with the amplitude asymmetry between the singing and silent wings arising from active neural suppression of the non-singing wing [52]. An alternative mechanism that would not require bilateral contraction of indirect flight muscles is mechanical coupling through the thorax. These hypotheses could be distinguished with simultaneous EMG recordings from the indirect flight muscles during wing switches.

Beyond song structure, our multi-animal 3D tracking revealed previously undescribed 3D postural dynamics during courtship: males adopted a lower body posture during running while singing (compared to running alone) and modulated their pitch angle to align vertically with the female’s position, a dimension of social orientation that is invisible to 2D tracking. These postural adjustments may reflect the competing demands of maintaining visual tracking of a moving target while controlling approach distance and body orientation [44, 50], and may direct acoustic flow of courtship song to be perceived by the female [54].

### 3.3 Advantages and limitations of our approach

Our multi-camera system achieved tracking accuracy that rivaled manual annotation, but several limitations constrain its current applications. First, the behavioral chamber (24 × 5.5 × 9.5 mm) restricts observations to relatively short locomotor sequences within a confined corridor. Extended or more naturalistic behavioral tracking would require either larger arenas with additional cameras or automated systems that follow the animal [60, 61], both of which would require additional hardware. Second, while our 50-keypoint skeleton captures all major body segments including legs, wings, head, and thorax, it does not resolve fine structures such as individual tarsal segments or antennal segments. Resolving individual tarsal segments would reveal how flies grip and release the substrate during each step, while tracking antennal movements would provide insight into how flies use active sensation to measure wind direction and intensity [62]. Tracking these structures would require higher magnification optics, which would further reduce the already small field of view. Third, our approach reconstructs detailed kinematics but does not directly measure dynamic variables like ground reaction forces or muscle forces, so biomechanical models trained from these data still do not have access to ground truth dynamics. Integration of the rainbow rig with substrate-embedded force sensors or inertial measurements [38] could address this gap in future iterations. The current system used white LEDs strobed synchronously with the camera exposure, which served the dual purpose of illuminating the arena and providing the broadband light needed to activate CsChrimson in our optogenetic experiments. However, this coupling between illumination and optogenetic stimulus limits experimental flexibility. In the future, using pulsed infrared illumination for imaging would free the visible spectrum for independent control of optogenetic stimulation intensity and timing, or for the presentation of controlled visual stimuli to the fly.

Despite these limitations, our system provides substantial advantages over existing approaches. Unlike tethered preparations, which constrain posture, alter ground reaction forces, and restrict the fly’s behavioral repertoire, our system captures natural kinematic variability across the full range of spontaneous behaviors of flies on the ground. Compared to existing 3D tracking systems optimized for larger animals, our telecentric lens configuration and custom calibration procedure are specifically designed to handle the optical challenges of millimeter-scale, high-speed imaging. The open-source publication of our hardware designs, calibration procedures, and trained JARVIS models will enable other laboratories to implement and extend this approach with minimal barriers to entry.

### 3.4 Future directions and broader applications

The methods we developed are immediately applicable to other small terrestrial arthropods where 3D kinematics have been difficult to obtain. Mosquitoes, ants, spiders, and other arthropods of similar size present comparable technical challenges, and our 3D pose tracking framework could be adapted with minimal modification. Beyond arthropods, the principles underlying our calibration and multi-camera triangulation approach generalize to any application requiring high-precision 3D tracking at millimeter scales.

The high-fidelity 3D kinematic dataset we generated fills a critical gap in neuromechanical modeling of *Drosophila* behavior. Both existing whole-body fly models (NeuroMechFly [27], Janelia Fly Body model [25]) have relied primarily on 2D kinematic data or 3D recordings from tethered animals. Our data reveal kinematic features that are particularly valuable for model training and validation: the 3D structure of joint rotations during turning, the running-like CoM dynamics that constrain leg spring mechanics, and the natural variability in inter-leg coordination at low speeds. These observations will enable more realistic imitation learning and provide ground truth for validating model predictions. As neuromechanical models mature toward incorporating connectome-derived circuit architectures, the demand for high-fidelity freely behaving kinematic data will only grow. Our open-source tools and dataset are designed to meet this need. Our demonstrations of pose estimation during grooming, optogenetics, and locomotion on complex terrain suggest a clear path toward collecting future datasets that systematically link identified neural circuit manipulations to their full-body biomechanical consequences.

## Supporting information

vid s1

vid s2

vid s3

vid s4

vid s5

vid s6

## Acknowledgments

We thank David Labonte, Simon Sponberg, Sama Ahmed, and members of the Tuthill and Brunton labs for discussions and helpful comments on the manuscript. We thank Andrea Gugiu (Janelia Experimental Technology) for contributing to Saturn IV LED fabrication; Grant Chou for developing a version of the PCA analysis on tethered locomotion kinematics and sharing his results and code. C.M. and T.T. are members of Janelia’s PTR-Bioimage Analysis team led by Sanna Koskela. We thank the Janelia Invertebrate Shared Resource team (Viruthika Vallanadu, Claire Angstadt, and Amanda Cavallaro) for fly husbandry support. This work was funded by the Howard Hughes Medical Institute (to R.E.J.), National Institutes of Health grants R01NS102333, R01NS128785, the New York Stem Cell Foundation, and a Pew Biomedical Scholar Award to J.C.T.; NIH grant R01NS145438 to J.C.T. and B.W.B.; NIH grants U01NS136507, R01NS136988, and the Richard & Joan Komen University Chair to B.W.B. J.C.T. is a New York Stem Cell Foundation – Robertson Investigator.

## Author Contributions

J.I.I. and E.T.T.A. performed data collection, curation, formal analysis, investigation, methodology, software development, validation, visualization, and writing. J.Y. developed the video acquisition, annotation, and 3D-pose estimation pipeline used in this study and contributed to investigation, software development, and validation. R.O. developed the fly rig camera calibration routine and integrated telecentric lens models into *red* and JARVIS. S.S. invented and developed the Saturn IV LED strobe light array used in this study, and contributed to investigation. F.A. contributed to investigation and methodology, including developing the first wing tracking model for courtship song detection. H.M.S. contributed to investigation and methodology, including prototyping fly chambers and planning and executing courtship experiments, and mentored F.A. N.R.M., J.W., T.T., and C.M. contributed to data curation, including manual annotation of thousands of 3D keypoint frames. W.C. developed and integrated a re-synchronization routine into the pose estimation software. J.V. co-developed the Saturn IV LED strobe light array and contributed to methodology. D.L.S. contributed to funding acquisition and mentorship of F.A. and H.M.S. R.E.J. contributed to methodology and designed and built the physical rainbow rig. B.W.B., J.C.T., and R.E.J. contributed to conceptualization, funding acquisition, supervision of the overall project, validation, visualization, and writing the manuscript.

## Data and Code Availability

All code developed for and used in this study is available on GitHub (https://github.com/elliottabe/3d_tracking_ik and https://github.com/moments-behavior). A curated 3D kinematics dataset can be downloaded at (https://bit.ly/3UWoXBF).

## Methods

### Rainbow rig design

To observe freely moving flies with sufficient spatiotemporal resolution, depth of field, and view sampling to enable robust 3D tracking, we used 7 high-speed cameras with telecentric lenses arranged in a semi-circle and aimed at the center of the fly arena floor. Cameras were synchronized with Precision Time Protocol (PTP) and strobe LED exposures were synchronized with camera shutters through camera GPIO hardware triggers. Below is a list of key hardware components and specifications.

### Lighting

Custom white strobe lights were designed and fabricated by Janelia Experimental Technology (https://www.janelia.org/support-team/janelia-experimental-technology). Each light module (named the Saturn IV Light Array) has 4 large white LED chips, a 30,000 *µ*F capacitor, and a 56 V power supply. We estimated one Saturn IV Light Array produces 75,000 lux during ON-time (measured at *∼*1 meter). In the rainbow rig, we positioned 2 Saturn IV Arrays to directly illuminate the fly arena, generating roughly 150,000 lux during image acquisition but just 6,000 lux perceptually (4% ON duty cycle). Diffuse strobe light also reflected off the camera mount plate to provide reasonably even multi-directional arena illumination. We measured the rise-time from darkness to maximum brightness to be *∼*2–3 *µ*s.

### Software tools for video acquisition, camera calibration, and keypoint annotation

All code and software used to acquire, calibrate, and annotate multi-camera video is available on GitHub [30, 31].

### Video acquisition

Videos were recorded using multi-camera streaming and acquisition software (named *orange* [30]). This software integrates APIs from Emergent Vision Technologies and NVIDIA to perform real-time H.264 video compression for up to eight 25GigE cameras at or near their maximum data rates on a single Threadripper PRO workstation. Video streams are synchronized with Precision Time Protocol (PTP) and routed from 25GigE network interface cards to GPU encoders for real-time compression using NVIDIA GPUDirect and NVENC. Two NVIDIA A16 GPUs in this workstation allow each camera to stream data to its own NVENC chip to enable continuous long-term recording of synchronized high-speed videos.

### Camera calibration

We developed a calibration routine that adapts the Direct Linear Transformation (DLTdv) method [63] to enable calibration for sets of cameras with telecentric lenses. This method requires a 3D calibration object, so we 3D printed an object with 23 small landmarks at precise 3D locations. We placed this object into the fly arena beneath the coverslips, so our calibration captured image shift induced by refraction through the glass arena walls. We imaged and annotated the pixel locations of every landmark in every camera view and performed an optimization to solve for each camera matrix. We integrated this DLT calibration routine for telecentric lenses into *red* annotation software.

To characterize the tracking accuracy of our rig, we calculated its reprojection error, which measures the distance in 2D camera image space from the labeled pixel location to the reprojected pixel location (after triangulation and reprojection). The per-landmark reprojection error for an example calibration was 0.48 *±* 0.26 pixels (5.9 *±* 3.2 *µ*m; *mean ± s.d.*) for N=272 landmarks (note this was computed on calibration training data). All cameras had a field of view of 1936 x 448 pixels with *∼*12.3 µm/pixel spatial resolution. To evaluate calibration quality on fly body parts, we labeled front left and front right tarsal tips of 51 frames in all 7 camera views and computed reprojection error of 1.48 *±* 0.69 pixels (18.2 *±* 8.5 *µ*m; *mean ± s.d.*).

### 3D keypoint annotation

Fly keypoints were labeled using multi-video streaming and annotation software (named *red* [31]). This application decodes synchronized videos using the NVIDIA NVDEC API, allowing users to view and label frames within large video datasets. Annotators identify frames of interest and then label each keypoint in two or more camera views before triangulating and reprojecting keypoint locations to all views. Images and keypoint locations are then exported for training a 3D pose estimation model (see Keypoint prediction below).

### Fly husbandry and behavioral experiments

For fly running and courtship experiments, we used wild-type Canton-S flies prepared by the Janelia Invertebrate Shared Resource. Flies were raised in standard cornmeal and molasses medium vials at 25C. Males and females were cold-plate sorted and housed separately in groups of *∼*20 individuals per vial. For single-fly running experiments, male and female flies aged 2-5 days were used. For courtship experiments, males were aged 4-5 days and females were aged 1-2 days to promote extended courtship displays without mating. For most running and courtship recordings, flies were transferred from culture vials to the chamber with an aspirator. For some running recordings, F1 progeny from a cross of Canton-S and empty stable split-Gal4 were used and were cold-anesthetized and transferred to the chamber with an aspirator. These flies were allowed to recover for 10 minutes before recording. Most recordings were 10-15 minutes long.

For optogenetic activation of running command neurons, a split-Gal4 line labeling DNg100 (w[1118]/ w[1118]; VT058557-GAL4.AD/+; R85F12-GAL4.DBD) was crossed with UAS-CsChrimson (*20xUAS-CsChrimson-tdTomato*) and progeny were aged for 14 days to allow for CsChrimson accumulation [57]. One day prior to recording, these flies were transferred to fly food containing all-trans retinal (1/250). Immediately before recording, flies were cold-anesthetized, decapitated, transferred to the recording chamber with an aspirator, and allowed to recover for 10 minutes before data collection. A red LED flashlight was used to induce neuronal activation. The warm red glow of this light can be seen in the behavioral videos (**Supplementary Video 5**).

### Keypoint prediction

We designed a 50-keypoint skeleton featuring landmarks across the head (midline point between antennae and dorso-posterior tip of each eye), thorax (scutellum tip, left and right wing hinges), abdomen (fourth abdominal stripe and abdomen tip), legs (forelegs coxa-thorax joints and every leg’s coxa-femur, femur-tibia and tibia-tarsus joint, plus the first, third and fifth tarsal segments) and wings (edge of the wing at vein 12 and 13). We annotated datasets to train JARVIS-HybridNet [32] models. JARVIS is a hybrid 2D/3D CNN that performs 3D pose estimation by back-projecting per-view 2D feature maps into a shared voxel grid and then applies 3D convolutions to localize keypoints in world space [64] together with per-keypoint confidence values. Approximately 2,500 frame-sets were labeled, each containing the same time point captured across the 7 cameras. Frames were chosen from three running video recordings and one of the courtship videos, selected for presenting unique poses across the chamber. To optimize labeling, intermediate models with fewer training frames were created and used to generate predictions on the data. Frames in which predictions showed the largest errors were relabeled and added to the training dataset for the next iteration, as these failure cases represent the most informative frames for improving model performance. Two JARVIS models were trained: A ’running-model’, containing labels from females and males performing running, grooming and courtship behaviors; a ’multi-animal courtship model’ containing courtship frames where both male and female flies were labeled. The ’running model’ was used to analyze both males and female flies during running and grooming and the ’multi-animal courtship model’ combined the single animal labels with simultaneous multi-animal labels and was used to analyze male flies during courtship, as well as perform a broad segmentation of females.

### Behavioral segmentation

#### Running data

A series of consecutive heuristic filters was created to extract running bouts from the JARVIS predictions. These consisted of 1) an 85 percent confidence threshold that eliminates poor predictions, 2) an upright filter, where all leg points must be in a lower Z position than the thorax, 3) a speed threshold, where the scutellum dX/dt should be higher than 5 mm/s, 4) a Y threshold for the tip of the legs that cleared out bouts where the fly legs get deflected by the glass walls, and 5) an X threshold to clear out climbing/demounting episodes at the start and end of the chamber.

#### Grooming data

A set of the same filters was used, removing the speed requirement and adding a Z-threshold for the forelegs (*Z >* 5 percentile for both legs at the same time). Antennal grooming was identified by the classic midline crossing of the forelegs during antennal rubbing, eye cleaning was identified by both tibia-tarsus joints being within 0.45 mm of its ipsilateral eye and proboscis/ocelli grooming by episodes where the head roll surpassed 35 degrees. Leg rubbing was identified by both tibia-tarsus joints being below the antennal base and less than 0.5 mm apart from each other in the XY plane.

#### Courtship data

A set of the running filters was used, removing the speed requirement (courtship sine usually happens during immobility) and adding a threshold for wing tip dZ/dt that ensures capturing song bouts. Wing tip position is calculated using the V13 keypoint. To detect sine and pulse within the singing bouts, we adapted the approach from FlySongSegmenter [45] to kinematic data. Pulse song was detected in the 200–380 Hz frequency band using a 4th-order zero-phase Butterworth bandpass filter. A smooth amplitude envelope was then obtained via the Hilbert transform, followed by low-pass filtering at 25 Hz. A noise floor was estimated from the bandpassed signal, and candidate pulses were identified as peaks in the envelope exceeding a fixed multiple of this noise floor, with a minimum inter-peak distance of 15 ms. Isolated peaks with no neighbor within 120 ms were discarded, as were trains whose median inter-pulse interval exceeded 80 ms. When multiple peaks were detected within a 10 ms window, only the one with the largest amplitude was kept. Sine song was detected using a multitaper spectral F-test, as implemented in FlySongSegmenter. Before analysis, regions surrounding detected pulse centers were masked out, and the masked signal was bandpass-filtered to 80–200 Hz. A sliding 100 ms window with a 10 ms step was applied, and at each position DPSS tapers were used to test whether the amplitude at any frequency in the 90–175 Hz range was better explained by a pure sinusoid than by the smooth broadband background. Windows that did not exceed the noise floor or fell within pulse train intervals were discarded. Consecutive significant windows with a consistent dominant frequency (within ±20%) were chained into candidate sine segments, and segments shorter than 80 ms were discarded.

All bouts were examined to ensure they contained only the target behavior and were written into summary text files containing start and end frames for each bout. Individual pulses were classified as slow-mode (*P_slow_*) or fast-mode (*P_fast_*) following [44]. For each detected pulse, a 25 ms window (±12.5 ms, 21 samples at 800 Hz) was extracted from the dominant wing tip Z-position trace, linearly detrended, z-scored, and sign-aligned so that the center sample was non-negative. Three features were computed from each aligned waveform: a symmetry index (cosine similarity between the first half and the time-reversed second half), a carrier frequency (energy-weighted mean of spectral bins exceeding 1/e of the peak magnitude), and the inter-pulse interval. Pulse type classification was performed once on all pulses pooled across fly pairs and wings. Waveforms were projected onto six principal components, and a two-component Gaussian mixture model with full covariance was fit to the resulting scores. The two clusters were assigned to pulse types by their mean symmetry index: the higher-symmetry cluster was designated *P_slow_* and the lower *P_fast_*, consistent with Clemens et al. A fixed random seed ensured deterministic assignments across runs. Each pulse was then labeled by projecting its waveform into the same PCA basis and assigning it to a cluster via the fitted GMM.

### Inverse kinematics

The 3D data exported by JARVIS were preprocessed so that keypoint names and order matched the tracking site order defined in the fly body model MuJoCo XML anatomy file. A Procrustes scale alignment was then computed from the mean pose (excluding the wings) across all frames in each clip and applied uniformly. A per-frame mask was then computed based on keypoint confidence (threshold was set at 0.5). We then applied bone length outlier constraints, as in Anipose [21]. For multi-animal tracking, we also imposed a centroid-jumping mask to prevent identity switching.

Once the data were preprocessed, we used a custom implementation of STAC-mjx [26] to run a two-phase optimization against the MuJoCo biomechanical body model: first fitting 3D marker attachment offsets on the rigid bodies to minimize reprojection error over an initial set of frames, then holding those offsets fixed and solving for full joint configurations (qpos) at every frame via inverse kinematics. The output of the solver produces the full kinematic and dynamic information (position, velocity, etc.) of the flies during the behaviors.

### Multi-animal Tracking

We built a custom framework to enable simultaneous 3D pose estimation of multiple freely behaving animals from synchronized multi-camera recordings. This pipeline utilizes an EfficientNet-based heatmap network (CenterDetect) to localize each instance of the fruit fly body centroid in each camera view. Around each center (one per fly) a 448 × 448 window is cropped from every camera, and the seven crops are lifted jointly by a query-based multi-view transformer. A pretrained DINOv3 ViT-B/16 backbone [65] (fine-tuned on our dataset), encodes each crop as a 28 × 28 grid of 768-dimensional patch tokens. Each token is made geometry-aware by back-projecting its own pixel through the calibrated camera to the line of sight along which its evidence lies. Our extension preserves this two-stage architecture but adds multi-instance detection, cross-camera identity resolution, and temporal tracking.

Multi-instance detection was achieved using a CenterDetect model trained on multi-animal data whose heatmaps contain one mode per individual, from which top-k peaks were extracted via iterative circular non-maximum suppression. Eight cross-attention layers read the fused tokens and predict, per query, a 3D position relative to the window center, a confidence, and per-slot existence and sex logits; four lighter layers, conditioned additionally on each camera’s projection parameters, decode the same queries into per-camera 2D positions and per-view visibilities.

We also integrated SAM3 (Segment Anything 3 [66]), a text-promptable segmentation model, as a detection and masking front-end. When applied per-frame, SAM3 was prompted with the label “insect” and returned pixel-accurate instance masks and centroids for each visible individual, thus substantially improving robustness when animals were touching or occluding one another. For bout-level processing, SAM3’s video propagation mode seeded identities on the first frame and tracked them forward with a memory bank; identities were then globally resolved across cameras by selecting the assignment that minimized multi-view reprojection error at an anchor frame where all animals were clearly visible. Finally, per-frame 3D body center estimates were linked across time using a multi-animal tracker with Hungarian matching and an identity-hold mechanism during brief occlusions.

### Swing/stance classification

The speed of the tip of each leg was used to assign a position in the step phase to each time point. This signal was chosen since there is a stark difference in speed between legs during stance, when the leg tip is stationary on the ground, and swing, when the leg moves forward. The speed was calculated by finite difference of the XYZ coordinates of the leg tip and Gaussian-smoothed (*σ* = 6 frames). The Hilbert transform was then applied to the mean-subtracted speed signal to obtain a continuous phase from *−π* to *π*. Phase = 0 corresponds to mid-swing and *|*phase*|* = *π* corresponds to mid-stance. A fixed threshold was then used to define swing and stance for each leg: frames with *|*phase*|< π/*2 were classified as swing and frames with *|*phase*| ≥ π/*2 as stance. Step cycles were arbitrarily defined by consecutive left foreleg swing onsets. A mean speed was assigned to each step cycle by averaging the forward speed across its frames.

### Inter-leg coordination across speeds

All legs were paired to L1 for comparison. For each leg pair the pooled distribution of relative phase was rendered as a von Mises kernel density estimate (*κ* = 12), with a band of *±*1 SD across flies, and the slow and fast distributions were overlaid on common axes. Mean direction and mean resultant length *R*were computed per fly and pooled. Distributions were compared with a two-sample Kuiper test, which is distribution-free and invariant to the choice of origin on the circle, so it does not depend on where the phase axis is cut. Asymptotic *p*-values were Bonferroni-corrected across the five panels.

### Joint-speed correlation analysis

The range of motion (RoM) of each joint was determined per step cycle by taking the difference between the maximum and minimum joint angle within that cycle, expressed in degrees. For each joint, the RoM values were averaged for both front (L1/R1), middle (L2/R2) and rear (L3/R3) legs to yield a bilateral estimate per cycle. For each fly, a Pearson correlation coefficient was computed between per-cycle RoM and per-cycle mean speed, then transformed into a Fisher z-score, and averaged across flies before back-transforming to a mean correlation coefficient. One-sample T-test was performed on the per-fly z-scores against the null hypothesis of zero mean correlation.

### PCA analysis of joint kinematics

Principal component analysis was used to reduce the dimensionality of the dataset and summarize joint kinematics. A feature matrix was constructed from the instantaneous joint angles and angular velocities of all joints across the 6 legs, z-scored, and decomposed into 10 principal components. Frames were divided into speed tertiles (low, medium and high) using the 33rd and 67th percentiles of step-cycle mean speed as boundaries. For each speed tertile, trajectories through the PC1–PC2 space were plotted and colored by the absolute phase offset of each leg from L1, with 0 indicating synchrony and *π* indicating antiphase. For the study of optogenetically induced locomotion, the feature matrix was mean centered using the healthy mean for each feature, and then projected into the healthy PCA space. Densities were estimated by Gaussian kernel density estimation with bandwidth 0.28 of the pooled standard deviation, evaluated on a common square 220 *×* 220 grid spanning the pooled data with 10% padding, and normalised to unit integral. Density is therefore expressed in cycles per PC^2^ and is directly comparable between conditions, since both distributions integrate to one over the same plane. Color encodes this absolute density in bands of fixed width, the width chosen automatically as 1, 2, 2.5 or 5 times a power of ten so as to give approximately nine bands up to the larger of the two peak densities (here 0.01 cycles/PC^2^, eight bands to 0.08). Every band edge is ruled with a thin line.

### Running dynamics analysis

#### Polar plots

The thorax height signal was detrended by subtracting a 100 ms rolling mean, isolating stride-coupled oscillations from slow postural drift; forward speed was used as is, since subtracting a per-cycle constant cannot displace a local maximum. A prominence-based peak finder was used to detect peaks in each signal (minimum prominence 9.8 *µ*m for height and 1.5 mm/s for speed, minimum separation 15 frames), which were then assigned to their corresponding step cycle. The phase of the left foreleg at the time of each peak was recorded. Step cycles were divided into speed tertiles as described above, and for each tertile, a polar histogram was constructed from the distribution of peak phases. Peaks were separated into a swing-half and a stance-half lobe at *±π/*2; within each lobe an arrow marks the phase shared by the two signals, its length giving the mean resultant of the paired height-minus-speed difference, and circles were used to mark the mean peak phase within each fly. Because peaks were detected independently for each signal, the total *n* differed slightly between speed and height peaks within each tertile.

#### Amplitude vs. speed

The amplitude of the height and speed signals was computed per step cycle, by calculating the root-mean-square fluctuation of each signal around its per-cycle mean. Per cycle amplitudes were plotted against step-cycle mean speed, with outliers beyond the 1st and 99th percentiles excluded. Binned means and standard deviations were overlaid using eight equal-count speed bins, and a linear regression was fitted.

### Coding agents

Anthropic’s large language model Claude Opus 5 was used as a coding agent in writing analysis code and making minor text corrections. Every output of the language model was verified by the authors.

## Supplementary Materials

**Supplementary Video 1:** An example running bout with superimposed keypoint tracking. Video is acquired at 800 Hz and played back at 80 Hz (10x slower).

**Supplementary Video 2:** An example bout of male courtship song with superimposed multi-animal keypoint tracking. Video playback is 13x slower than realtime.

**Supplementary Video 3:** Replay of courtship behavior, visualized through inverse kinematics onto biomechanical fly body models.

**Supplementary Video 4:** An example bout of anterior grooming, played back 10x slower than realtime.

**Supplementary Video 5:** An example locomotion bout driven by optogenetic activation of DNg100 (real time playback). The fly speeds up when the red light turns on 0.7 seconds into the video.

**Supplementary Video 6:** An example of a fly traversing a 3D-printed complex terrain of a series of steps (played back 10x slower).

---

**Supplementary Figure 1:**
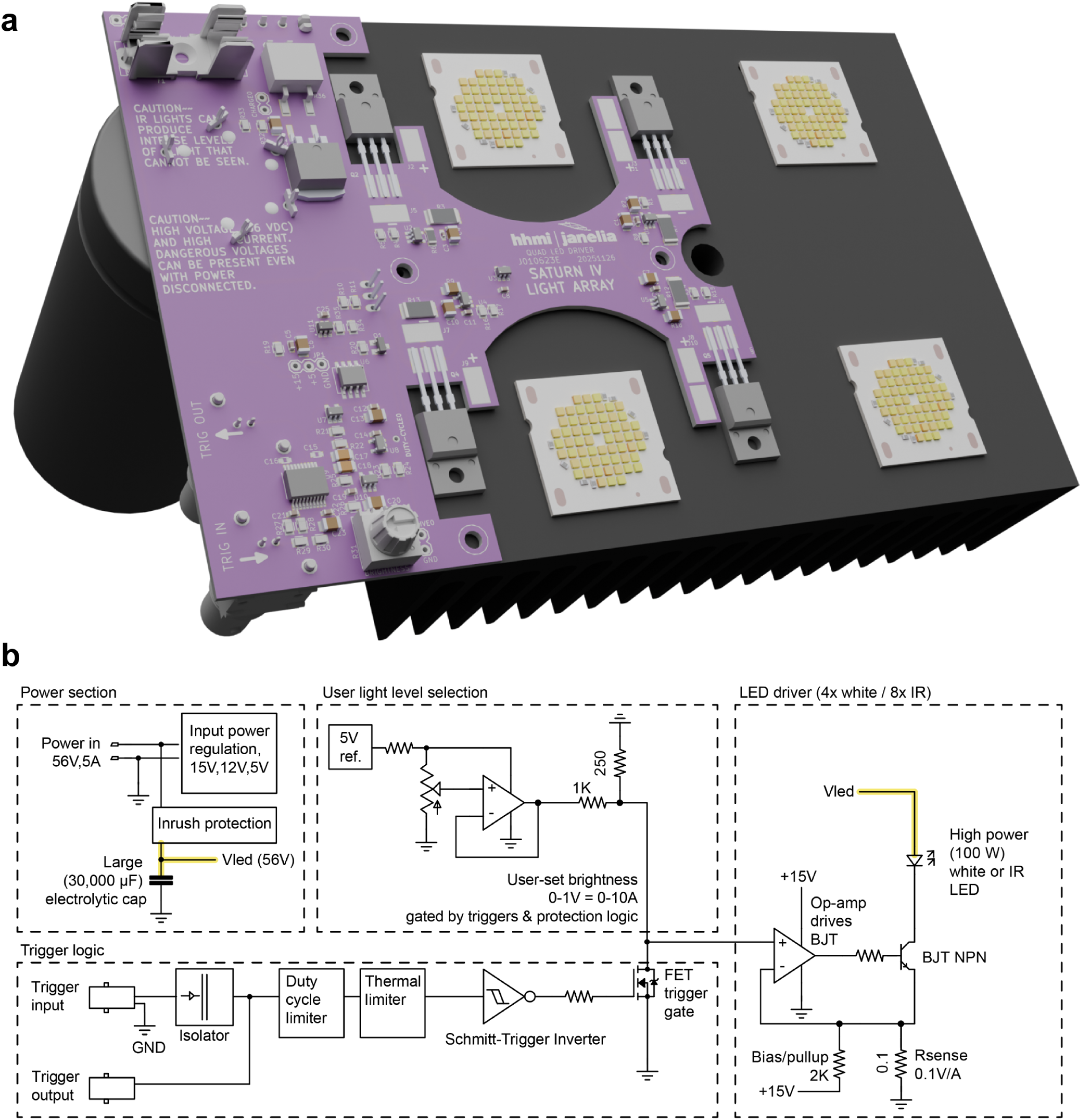
Saturn IV Light Array. **a**, CAD model of the Saturn IV Light Array. **b**, Simplified block diagram of the strobe lights: Input power (56 V, 250 W) is used to supply the LED power as well as the auxiliary supply rails. A large bulk capacitance on the 56 V LED power (*V*_led_) supports pulsed high-current operation without significant voltage drop. An inrush limiting circuit allows the capacitor to charge slowly at power on and keeps the LED drivers off until voltages are stable. A precision 5 V reference and user-adjustable potentiometer generate a 0– 1 V current-command signal, corresponding to 0–10 A LED current, to adjust brightness. This brightness command signal is then gated by a trigger input, typically driven by the camera’s exposure output signal. Flash duration is determined by the width of this trigger input. Trigger width is further gated by duty-cycle and thermal limiters. Each of the 4 LED channels (each can power one 100 W 30 V white or two 100 W 15 V IR LEDs in series) uses an op-amp-controlled NPN current sink, with feedback from a 0.1 Ω sense resistor providing a transconductance of 0.1 V/A. A trigger output enables daisy-chaining of strobes.

**Supplementary Figure 2:**
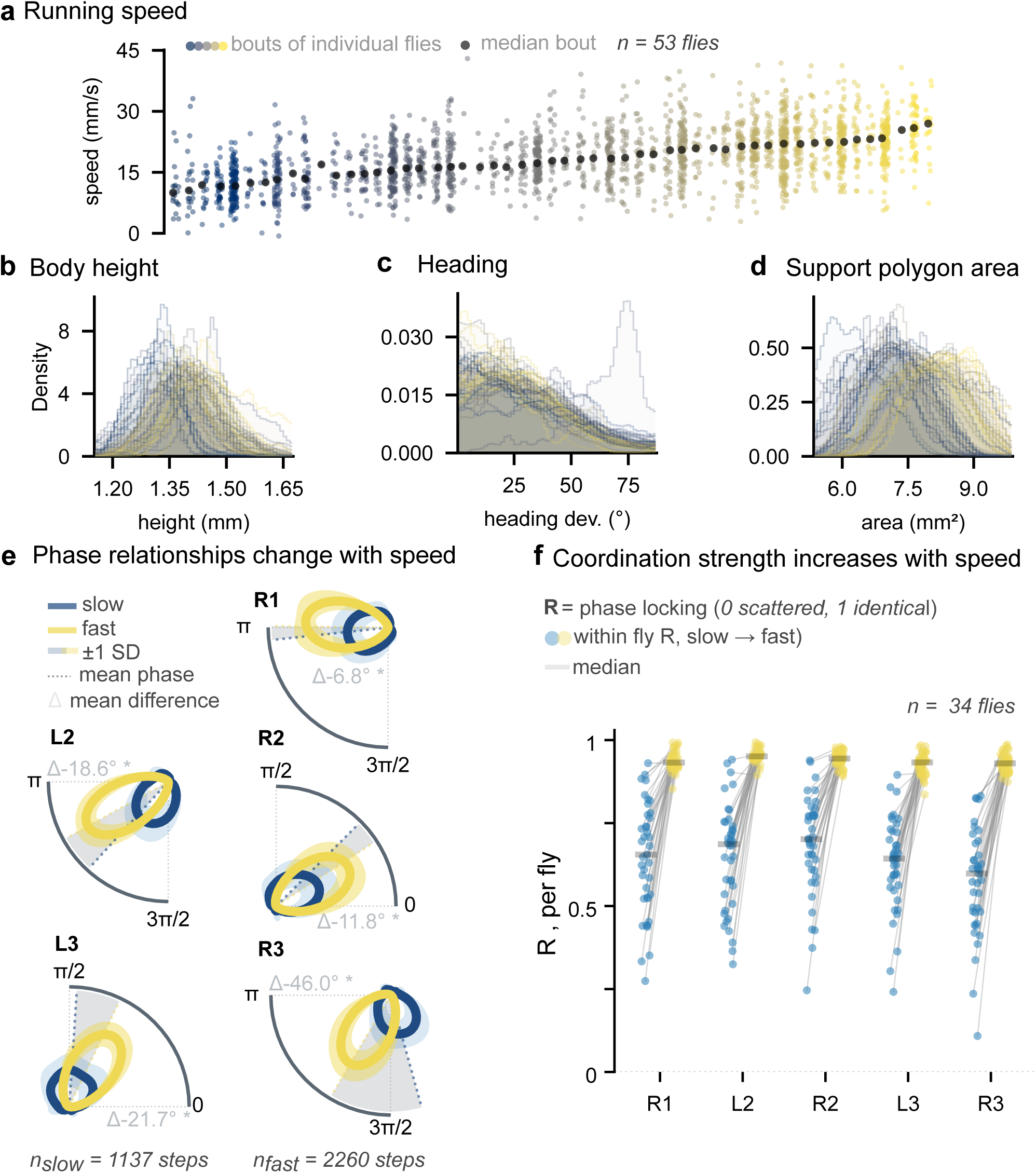
Running dataset overview. **a**, Per-bout running speed. Each column is one fly (n=53, 32 males and 21 females, single housed) ordered left to right by median speed. Each circle in a column corresponds to a running bout (n=2,213). The median forward speed across flies is 17.8 mm/s. **b-d**, Pooled distributions of (b) body height, measured as the difference between the Z component of the scutellum and the legs in stance for every frame, (c) heading, measured as the deviation from the corridor axis (0° = along the corridor, 90° = perpendicular; median 26.4°) and (d) support polygon area, measured as the area inside the polygon drawn by the XY component of the tip of the legs. Each fly contributes one transparent filled step histogram computed on shared bins (60 bins spanning the pooled 0.5–99.5th percentile, smoothed with a Gaussian of *σ* = 1.2 bins). **e**, Phase offsets to the foreleg across the speed range. Each panel represents a comparison of one of the legs to the left foreleg (L1). Curves are von Mises kernel density estimates (k=12) of the phase of that leg’s swing onset within the L1 stride cycle, averaged across flies and drawn for the slow and fast tertiles; shading is *±*1 SD across flies. Because tertiles are computed over frames, the two groups hold equal recording time rather than equal cycle counts, so group sizes differ by design (n = 1,050–1,168 slow vs. 2,154–2,191 fast phase comparisons per panel). Statistics are a two-sample Kuiper test per comparison, with *V* = *D*^+^ + *D^−^* as the effect size and p Bonferroni-corrected for the five panels; V = 0.30–0.57, all *p <* 0.01 (reported as *). **f**, Coordination strength per fly. Each dot is one fly’s mean resultant length R of the phase offsets for that leg, slow (left, blue) and fast (right, amber); horizontal bars are group medians and hairlines join the same fly across groups. Median R rises with speed for every leg (R1 0.70 to 0.93, L2 0.72 to 0.95, R2 0.73 to 0.95, L3 0.69 to 0.93, R3 0.63 to 0.93; Δ*R* = +0.22 to +0.30, 33–35 paired flies per leg comparison).

